# CAR-T cells targeting CCR9 and CD1a for the treatment of T cell acute lymphoblastic leukemia

**DOI:** 10.1101/2024.09.02.610843

**Authors:** Néstor Tirado, María José Mansilla, Alba Martínez-Moreno, Juan Alcain, Marina García-Peydró, Heleia Roca-Ho, Narcis Fernandez-Fuentes, Alba Garcia-Perez, Mercedes Guerrero-Murillo, Aïda Falgàs, Talia Velasco-Hernandez, Meritxell Vinyoles, Clara Bueno, Pablo Engel, E Azucena González, Binje Vick, Irmela Jeremias, Aurélie Caye-Eude, André Baruchel, Hélène Cavé, Eulàlia Genescà, Jordi Ribera, Marina Díaz-Beyá, Manuel Ramírez-Orellana, Montserrat Torrebadell, Víctor M Díaz, María L Toribio, Diego Sánchez-Martínez, Pablo Menéndez

**Affiliations:** Josep Carreras Leukaemia Research Institute (IJC), Barcelona, Spain; Red Española de Terapias Avanzadas (TERAV), Instituto de Salud Carlos III, Madrid, Spain; OneChain Immunotherapeutics S.L, Barcelona, Spain; Centro de Biología Molecular Severo Ochoa CSIC-UAM, Madrid, Spain; Department of Biomedicine, School of Medicine, University of Barcelona, Barcelona, Spain; Centro de Investigación Biomédica en Red de Cáncer (CIBERONC), Madrid, Spain; Department of Immunology, Hospital Clínic de Barcelona, Barcelona, Spain; Research Unit Apoptosis in Hematopoietic Stem Cells, Helmholtz Munich, German Research Center for Environmental Health (HMGU), Munich, Germany; Department of Pediatrics, Dr. von Hauner Children’s Hospital, LMU University Hospital, LMU Munich, Germany; Department of Genetics, University Hospital Robert Debré, Paris, France; INSERM Institut de Recherche Saint Louis, Paris, France; Department of Hematology, Institut Català d’Oncologia-Hospital Germans Trias i Pujol, Badalona, Spain; Department of Hematology, Hospital Clínic de Barcelona, Barcelona, Spain; Department of Pediatric Hematology and Oncology, Hospital Infantil Universitario Niño Jesús, Universidad Autónoma de Madrid, Madrid, Spain; Institut de Recerca Pediàtrica Sant Joan de Déu, Barcelona, Spain; Department of Hematology, Hospital Sant Joan de Déu, Barcelona, Spain; Universitat Internacional de Catalunya, Sant Cugat del Vallès, Spain; Aragon Health Research Institute (IIS Aragón), Zaragoza, Spain; Aragon Foundation for Research & Development (ARAID), Zaragoza, Spain; Institució Catalana de Recerca i Estudis Avançats (ICREA), Barcelona, Spain

**Author notes:** **Correspondence should be addressed to** PM DS-M MLT.

## Abstract

T cell acute lymphoblastic leukemia (T-ALL) is an aggressive malignancy characterized by high rates of induction failure and relapse, and effective targeted immunotherapies are lacking. Despite promising clinical progress with genome-edited CD7-directed CAR-T cells, which present significant logistical and regulatory issues, CAR-T cell therapy in T-ALL remains challenging due to the shared antigen expression between malignant and healthy T cells. This can result in CAR-T cell fratricide, T cell aplasia, and the potential for blast contamination during CAR-T cell manufacturing. Recently, CAR-T cells have been described that target non-pan-T antigens, absent on healthy T cells but expressed on specific T-ALL subsets. These antigens include CD1a (NCT05679895), which is expressed in cortical T-ALL, and CCR9. We show that CCR9 is expressed on >70% of T-ALL patients (132/180) and is maintained at relapse, with a safe expression profile in healthy hematopoietic and non-hematopoietic tissues. Further analyses showed that dual targeting of CCR9 and CD1a could benefit ∼86% of patients with T-ALL, with a greater blast coverage than single CAR-T cell treatments. We therefore developed, characterized, and preclinically validated a novel humanized CCR9-specific CAR with robust and specific antileukemic activity as a monotherapy *in vitro* and *in vivo* against cell lines, primary T-ALL samples, and patient-derived xenografts. Importantly, CCR9/CD1a dual-targeting CAR-T cells showed higher efficacy than single-targeting CAR-T cells, particularly in T-ALL cases with phenotypically heterogeneous leukemic populations. Dual CCR9/CD1a CAR-T therapy may prevent T cell aplasia and obviate the need for allogeneic transplantation and regulatory-challenging genome engineering approaches in T-ALL.

## INTRODUCTION

T cell acute lymphoblastic leukemia (T-ALL) is a clonal hematologic malignancy characterized by a block in differentiation, resulting in the accumulation of T cell lineage lymphoblasts, and often presents with leukocytosis, cytopenias, and extramedullary infiltration^1^. It is a highly heterogeneous disease both phenotypically and genetically, with recurrent mutations in transcription factors and signaling pathways involved in hematopoietic homeostasis and T cell development^2–4^. T-ALL accounts for ∼15% and ∼25% of all pediatric and adult ALL cases, respectively. Treatment is based on intensive multi-agent cytotoxic chemotherapy^1^. Despite a cure rate of ∼80% in children^2,3^, long-term survival is <40% in adult patients who can tolerate intensive chemotherapy^4^. More than half of patients relapse or fail to respond to standard therapy, resulting in a very poor prognosis. Median overall survival for patients with relapsed/refractory (R/R) disease is ∼8 months^5^. For R/R T-ALL, the standard approach to achieve remission typically requires intensive re-induction chemotherapy followed by allogeneic hematopoietic stem cell transplantation (alloHSCT). However, this is associated with significant toxicity and high failure rates, highlighting the urgent need for new targeted and safe therapeutic strategies for patients with R/R T-ALL.

Unlike B cell malignancies, which have effective immunotherapy target antigens such as CD19, CD22, or CD20, the are currently no approved immunotherapies for T-ALL^6–9^. A major challenge in the development of immunotherapies against T cell malignancies is the lack of safe and actionable tumor-specific antigens^10,11^. The high phenotypic similarity between effector T cells and leukemic lymphoblasts not only complicates the manufacture of autologous CAR-T cells directed against pan-T antigens such as CD7 or CD5, but also induces fratricide and T cell aplasia^12–16^. Recent clinical studies have addressed these limitations by using either genome-edited or expression blocker-engineered CD7-directed CAR-T cells^17–22^. While elegant, these strategies present significant logistical and regulatory challenges, and are limited to the use of allogeneic effector T cells and to patients with a donor available for rescue therapy with alloHSCT.

A solution to these limitations is to direct CAR-T cells against non-pan-T antigens, which are expressed on blasts but not on healthy T lymphocytes. This strategy would facilitate the manufacture of autologous CAR-T cells while also avoiding both fratricide and immune toxicity^23–28^. In this context, we previously identified CD1a as an immunotherapeutic target for the treatment of cortical T-ALL with a safe profile in non-hematopoietic and hematopoietic tissues, including normal T cells. This allowed us to generate and validate CD1a-directed CAR-T cells, which are now being tested in a phase I clinical trial (NCT05679895)^25,29^. However, CD1a is expressed solely in cortical T-ALL cases, a subtype that accounts for only ∼30% of all T-ALL cases, while sparing other T-ALL subtypes that are critically associated with higher refractoriness and relapse rates.

Expression of the chemokine receptor CCR9, a G protein-coupled receptor for the ligand CCL25, has recently been suggested to be restricted to two-thirds of T-ALL cases, and CAR-T cells targeting CCR9 were resistant to fratricide and had potent antileukemic activity in preclinical studies^30,31,27^. Here, we show that dual targeting of CCR9 and CD1a may benefit a large proportion of T-ALL cases, with greater blast coverage than treatment with single-targeting CAR-T cells. We preclinically validate a novel humanized CCR9-specific CAR with robust and specific antileukemic activity as a monotherapy *in vitro* and *in vivo* and demonstrate the advantage of dual CCR9- and CD1a-targeting CAR-T cells, particularly in T-ALL cases with phenotypically heterogeneous leukemic populations. We propose a highly effective CAR-T cell strategy for T-ALL that may prevent T cell aplasia and circumvent the need for alloHSCT and regulatory-challenging genome engineering approaches.

## METHODS

### Donor and patient samples

Research involving human samples was approved by the Clinical Research Ethics Committee (HCB/2023/0078, Hospital Clínic, Barcelona, Spain). Thymus (*n*=4), peripheral blood (PB, *n*=18), and bone marrow (BM, *n*=13) samples were obtained from healthy individuals. BM samples were sourced from healthy donor transplantation leftovers and thymuses were obtained from patients undergoing thymectomy from thoracic surgeries. Diagnostic and relapse primary T-ALL samples (*n*=180) were obtained from the sample collections of the participating hospitals after informed consent (IGTP Biobank, PT17/0015/0045).

### Cell lines

MOLT4, SupT1, and MV4;11 cell lines were purchased from the DSMZ cell line bank and cultured in RPMI 1640 supplemented with 10% fetal bovine serum (FBS). CCR9-knockout (KO) and CD1a KO MOLT4 cells were generated by CRISPR-mediated genome editing. 500,000 cells were electroporated using the Neon Transfection System (Thermo Fisher Scientific) with a Cas9/crRNA:tracrRNA complex (IDT). A crRNA guide was designed for each gene: *CCR9* 5’-GAAGTTAACGTAGTCTTCCATGG-3’ and *CD1A* 5’-TATTCCGTATACGCACCATTCGG-3’. After electroporation, cells were recovered, and the different KO clones were FACS-sorted and purity confirmed (>99%).

### CCR9 monoclonal antibody and generation of a humanized scFv

Monoclonal antibodies (mAbs) reactive with human CCR9 were generated using hybridoma technology (ProteoGenix). Anti-CCR9 antibody-producing hybridomas were generated by immunizing mice with different peptides derived from the human extracellular N-terminal region of human CCR9 fused to a KLH carrier protein to improve stability and the immune response. After hybridoma subcloning and initial ELISA screenings, supernatants from individual clones were analyzed by flow cytometry for reactivity against CCR9-expressing MOLT4 and 300.19-hCCR9 cells and their respective negative controls (MOLT4 CCR9 KO and wild type 300.19 cells). One hybridoma (clone #115, IgG1 isotype) was selected, its productive IgG was sequenced, and the V_H_ and V_L_ regions were used to derive the murine single-chain variable fragment (scFv) using the Mouse IgG Library Primer Set (Progen), as described^25,32^.

For humanization, a sequence search was performed in the IMGT database^33^ to identify immunoglobulin genes with the highest identity to both the V_H_ and V_L_ domains of murine antibody #155. The highest murine-human sequence identities were *IGHV1-3*01* for V_H_ (60% identity) and *IGKV2D-29*02* for V_L_ (82% identity); the number of different residues (with different levels of conservation) excluding the complementarity-determining regions (CDRs) was 32 and 13, respectively. The CDRs, including the Vernier regions, were grafted onto the human scaffolds. A structural model of the murine scFv was used to identify other structurally important residues in the antibody that differed from the corresponding positions in the humanized versions and should be retained (e.g., buried residues, residues located at the interface, etc.). Two humanized candidates were generated, each with different degrees of residue substitution at non-conserved positions. The sequence-based humanized candidate #1 (H1) aimed to have only the changes necessary to transfer CDRs, Vernier, and structurally important residues to the human scaffold, while the humanized candidate #2 (H2) allowed for some additional changes to be made to better match the human sequence.

### CAR design and vectors, lentiviral production, and T cell transduction

Single scFvs (CD1a H and CCR9 M, H1, and H2), all possible configurations of tandem CAR constructs (*n*=8) and four different configurations of bicistronic CAR, were cloned into the clinically validated pCCL lentiviral backbone containing the human CD8 hinge and transmembrane (TM) domains, 4-1BB and CD3ζ endodomains, and a T2A-eGFP reporter cassette. All constructs contain the signal peptide (SP) derived from CD8α (SP1) upstream (5’) of the first scFv. Two different SPs derived from either human IgG1 (SP2) or murine IgG1 (SP3) were used for the second CAR in bicistronic constructs. Third-generation lentiviral vectors were generated in 293T cells by co-transfection of the different pCCL expression plasmids, pMD2.G (VSV-G) envelope, and pRSV-Rev and pMDLg/pRRE packaging plasmids using polyethylenimine (Polysciences)^34^. Viral particle-containing supernatants were collected at 48 and 72 h after transfection and concentrated by ultracentrifugation.

PB mononuclear cells were isolated from buffy coats of healthy donors by density-gradient centrifugation using Ficoll-Paque Plus (Merck). Buffy coats were sourced from the Catalan Blood and Tissue Bank. T cells were activated for 2 days in plates coated with anti-CD3 (OKT3) and anti-CD28 (CD28.2) antibodies (BD) and transduced with CAR-encoding lentiviral particles at a multiplicity of infection (MOI) of 10. T cells were expanded in RPMI 1640 medium containing 10% heat-inactivated FBS, penicillin-streptomycin, and 10 ng/mL interleukin (IL)-7 and IL-15 (Miltenyi Biotec). Expression of CAR molecules in T cells was detected by flow cytometry using the eGFP reporter signal and biotin-SP goat anti-mouse IgG, F(ab’)_2_ (Jackson ImmunoResearch) and PE-conjugated streptavidin (Thermo Fisher Scientific). Vector copy number (VCN) was determined by qPCR using Light Cycler 480 SYBRGreen I Master (Roche) using primers designed against the WPRE proviral sequence, as described^35,36^. Absolute quantification was used to determine VCN/genome, adjusted to the percentage of transduction of each CAR as determined by flow cytometry analysis.

### *In vitro* cytotoxicity and cytokine secretion assays

Target cells (100,000 to 300,000 cells/well) were labeled with eFluor 670 dye (Thermo Fisher Scientific) and incubated in a 96-well round bottom plate with untransduced or CAR-transduced T cells at the indicated effector:target (E:T) ratios for 24 h. Cytotoxicity was assessed by flow cytometric analysis of residual live target cells (eFluor 670^+^ 7-AAD^−^). For primary T-ALL blasts, the absolute number of live target cells was also determined using Trucount tubes (BD). Additional wells containing only target cells (“no effector”; NE) were always plated as controls. Transduction percentages were adjusted across conditions when comparing multiple CAR constructs. Quantification of the pro-inflammatory cytokines IFN-γ, TNF-α, and IL-2 was performed by ELISA using BD OptEIA Human ELISA kits (BD) on supernatants collected after 24 h of target cell exposure.

### Flow cytometry

The fluorochrome-conjugated antibodies against CD1a (HI149), CD3 (UCHT1), CD4 (SK3), CD7 (M-T701), CD8 (SK1 and RPA-T8), CD14 (MφP9), CD19 (HIB19), CD34 (8G12), CD38 (HIT2), CD45 (HI30, 2D1), HLA-ABC (G46-2.6), mouse IgG1, κ isotype control (X40) and the 7-AAD cell viability solution were purchased from BD. Antibodies against CCR9 (L053E8), TCRαβ (IP26), His tag (J095G46), and mouse IgG2a, κ isotype control (MOPC-173) were purchased from BioLegend. Antibodies against CD4 (13B8.2), CD34 (581), and TCRγδ (IMMU510) were purchased from Beckman Coulter. Alexa Fluor 647-conjugated anti-mouse IgG (H+L) was purchased from Cell Signaling Technology. Samples were stained with mAbs (30 min at 4°C in the dark) and erythrocytes were lysed (when applicable) using a FACS lysing solution (BD). Isotype-matched nonreactive fluorochrome-conjugated mAbs were used as controls. Cell acquisition was performed on a FACSCanto II and analyzed using FACSDiva and FlowJo v10 software (BD). All gating strategies and analyses are shown in **Fig. S1**.

### *In vivo* assessment of CAR-T cell efficacy in T-ALL models

*In vivo* experiments were performed in the Barcelona Biomedical Research Park (PRBB) animal facility. All procedures were approved by and performed according to the guidelines of the PRBB Animal Experimentation Ethics Committee in agreement with the Generalitat de Catalunya (DAAM11883). All mice were housed under specific pathogen-free conditions. Seven-to twelve-week-old NOD.Cg-*Prkdc^scid^ Il2rg^tm1Wjl^*/SzJ (NSG) mice (The Jackson Laboratory) were sublethally irradiated (2 Gy) and systemically transplanted with 1×10^6^ T-ALL patient-derived xenograft (PDX) cells via the tail vein. Two to three weeks later, PB and BM samples were collected to assess the leukemic burden and establish the different treatment groups prior to CAR-T cell injection (3-4×10^6^ cells) via the tail vein. For *in vivo* studies with MOLT4 cell lines, phenotypically heterogeneous MOLT4 cells (CCR9^+^CD1a^+^, CCR9^−^CD1a^+^, CCR9^+^CD1a^−^; 1:1:1 ratio, 1×10^6^ cells) were transplanted three days before CAR-T cell administration. Tumor burden was monitored weekly by flow cytometry of PB samples. For luciferase-expressing T-ALL models MOLT4 and ALL-843^37^, tumor growth was monitored weekly by bioluminescence measurement after intraperitoneal administration of 60 mg/kg of D-luciferin (PerkinElmer). Bioluminescence was evaluated using Living Image software (PerkinElmer). Mice were sacrificed when signs of disease were evident. Spleens were manually dissected, and a single-cell suspension was obtained using a 70 µm strainer. Samples were stained and processed for flow cytometry as described above.

### Statistical analysis

For CAR-T cell expansion, cytotoxicity, and *in vivo* studies, two-way ANOVA with Tukey’s multiple comparison adjustments was used to compare the different groups, with untransduced T cells serving as controls. For cytokine release assays, a one-way ANOVA with Dunnett’s multiple comparison adjustment was used. All statistical tests were performed using Prism 6 (GraphPad Software). The number of biological replicates is indicated in the figure legends. Significance was considered when *p*-values were lower than 0.05 (ns, not significant; *p<0.05; **p<0.01; ***p<0.001).

## RESULTS

### CCR9 is a safe and specific target for T-ALL

Apart from CD1a, there are no other non-pan-T safe markers available for targeting T cell tumors with adoptive cellular immunotherapy. Ideally, immunotherapy targets should be expressed in tumor cells but not in healthy tissues, including T cells. We first analyzed CCR9 expression by flow cytometry in a cohort of 180 T-ALL samples (**Fig. 1a**). Consistent with a previous report from the Great Ormond Street Hospital/University College London group^27^, we found that 73% (132/180) of T-ALL samples were CCR9^+^, with variable levels of expression (using a cut-off of ≥20% expression for positivity) (**Fig. 1a, S1a**). Importantly, non-leukemic CD4^+^ and CD8^+^ T cells from the same patients remained CCR9^−^. This proportion of CCR9 positivity in T-ALL was maintained (64-76%) when patients were stratified into different maturation subtypes according to the EGIL immunophenotypic classification^38^ (**Fig. 1b**). Notably, the proportion of CCR9^+^ T-ALL cases increased considerably in relapse samples (92%, 12/13), with a much higher blast coverage than observed at diagnosis (**Fig. 1c**).

**Figure 1.**
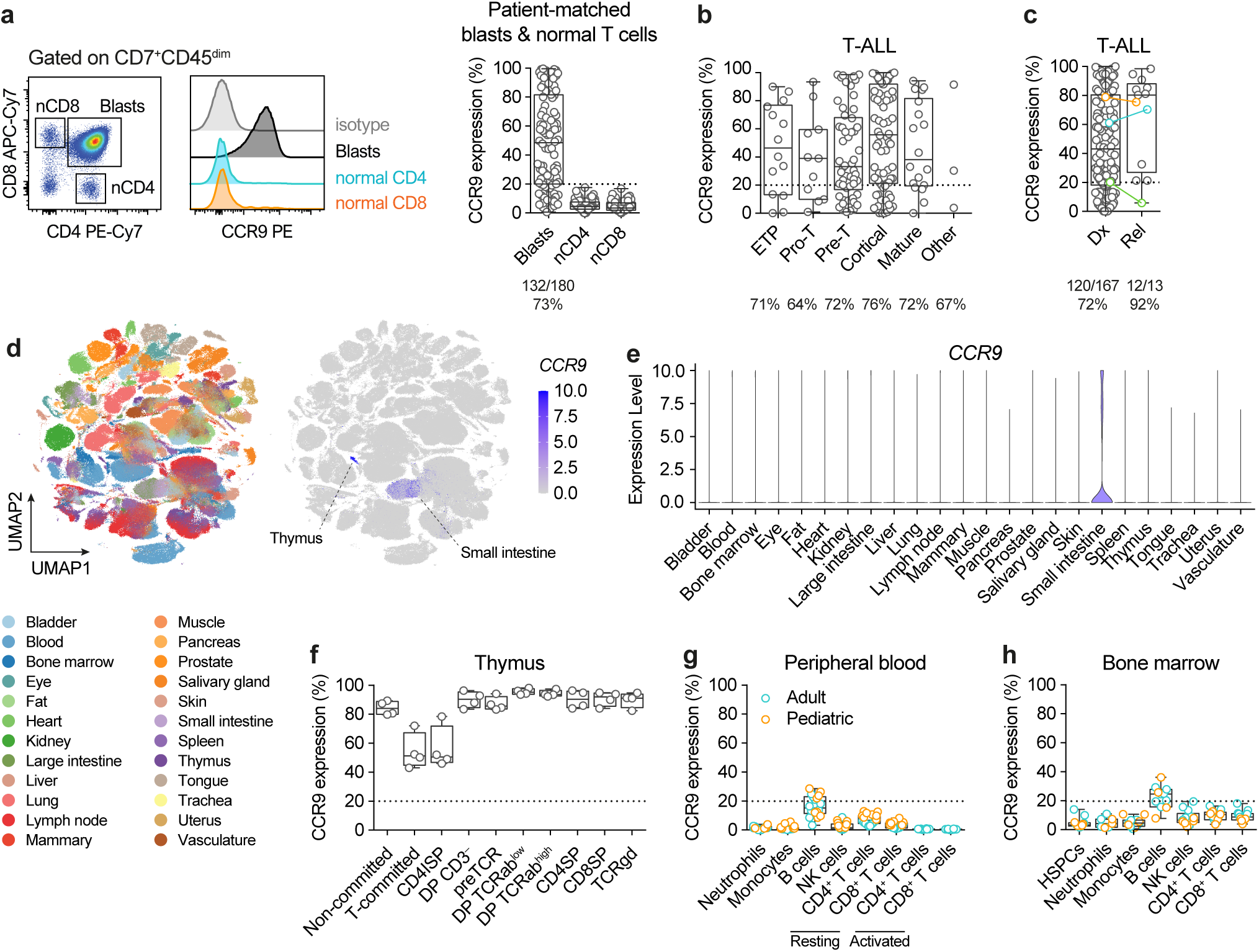
CCR9 is a safe and specific target for T-ALL. (**a**) Flow cytometry analysis of CCR9 in patient-matched leukemic blasts and normal (n) CD4^+^ and CD8^+^ T cells in 180 T-ALL patients. Left panels, representative flow cytometry plot. (**b,c**) Expression of CCR9 in the cohort, with T-ALL samples classified by developmental stage (**b**) or disease stage, at diagnosis (Dx) or relapse (Rel) (**c**). Three cases with Dx-Rel matched samples are color-coded. (**d**) UMAP representation showing organ/tissue annotation and *CCR9* expression levels in 483,152 cells from human healthy tissues (Tabula Sapiens scRNAseq dataset). (**e**) Violin plots for *CCR9* expression levels across tissues identified in (d). (**f-h**) CCR9 expression in the indicated leukocyte populations in relevant hematological tissues: thymus (**f**, *n*=4), PB (**g**, *n* =18), and BM (**h**, *n*=13).

An immunotherapeutic target must meet a stringent safety profile, ensuring that it is not expressed in other hematopoietic or non-hematopoietic cell types. To assess the safety profile of CCR9, we first examined its expression in the Tabula Sapiens single-cell RNAseq dataset, a human reference atlas comprising 24 different tissues and organs from adult healthy donors^39^. We found a complete absence of CCR9 expression in all tissues except for minor subsets of thymic cells and small intestinal resident lymphocytes (**Fig. 1d,e**). We confirmed the expression of CCR9 in all thymocyte subpopulations along T cell development by flow cytometric analysis of postnatal thymuses (*n*=4, **Fig. 1f, S1b**). However, the same analysis of healthy pediatric and adult PB (*n*=18) and BM (*n*=13) samples revealed that, with the exception of expression in 10-30% of B cells, CCR9 was minimally expressed in all major leukocyte subpopulations analyzed, including CD34^+^ hematopoietic stem/progenitor cells (HSPCs) and resting and CD3/CD28-activated T cells (**Fig. 1g,h; S1c,d**). In sum, CCR9 is expressed in a high proportion of T-ALL patients, particularly at relapse, while exhibiting a safety profile characterized by low or absent expression in healthy tissues, CD34^+^ HSPCs, and T lymphocytes. This highlights its potential as a target for the development of CCR9-directed, fratricide-resistant, safe CAR-T cell therapy.

### CCR9 CAR-T cells are highly effective against T-ALL blasts

Next, we sought to develop a humanized CCR9-directed CAR for the treatment of R/R T-ALL. The highly hydrophobic nature and insolubility of the CCR9 protein led us to use the CCR9 N-terminal extracellular tail for mouse immunization and subsequent generation of murine CCR9 antibody-producing hybridomas. After hybridoma subcloning and individual testing for CCR9 reactivity, one clone (#115) demonstrated specificity (**Fig. 2a**). The productive IgG was sequenced and the V_H_ and V_L_ regions were used to derive the murine scFv for CAR design. Two additional humanized scFvs were generated by structural fitting and modeling of the CDRs and neighboring regions into human IgG scaffolds (**Fig. S2a**). Epitope mapping using overlapping peptides from the CCR9 N-terminus revealed the CCR9 epitope recognized by the clone #115 scFv (**Fig. S2b**). The murine (M) and both humanized (H1 and H2) CCR9 scFvs were cloned into the clinically validated pCCL-based second-generation CAR lentiviral backbone, including a T2A-eGFP reporter cassette (**Fig. 2b**). Primary T cells were successfully transduced, and CAR expression detected by anti-F(ab’)_2_ staining, which correlated with the eGFP signal (**Fig. 2c**). All CCR9 CAR-T cells showed identical expansion to untransduced T cells, demonstrating the absence of fratricide (**Fig. 2d**).

**Figure 2.**
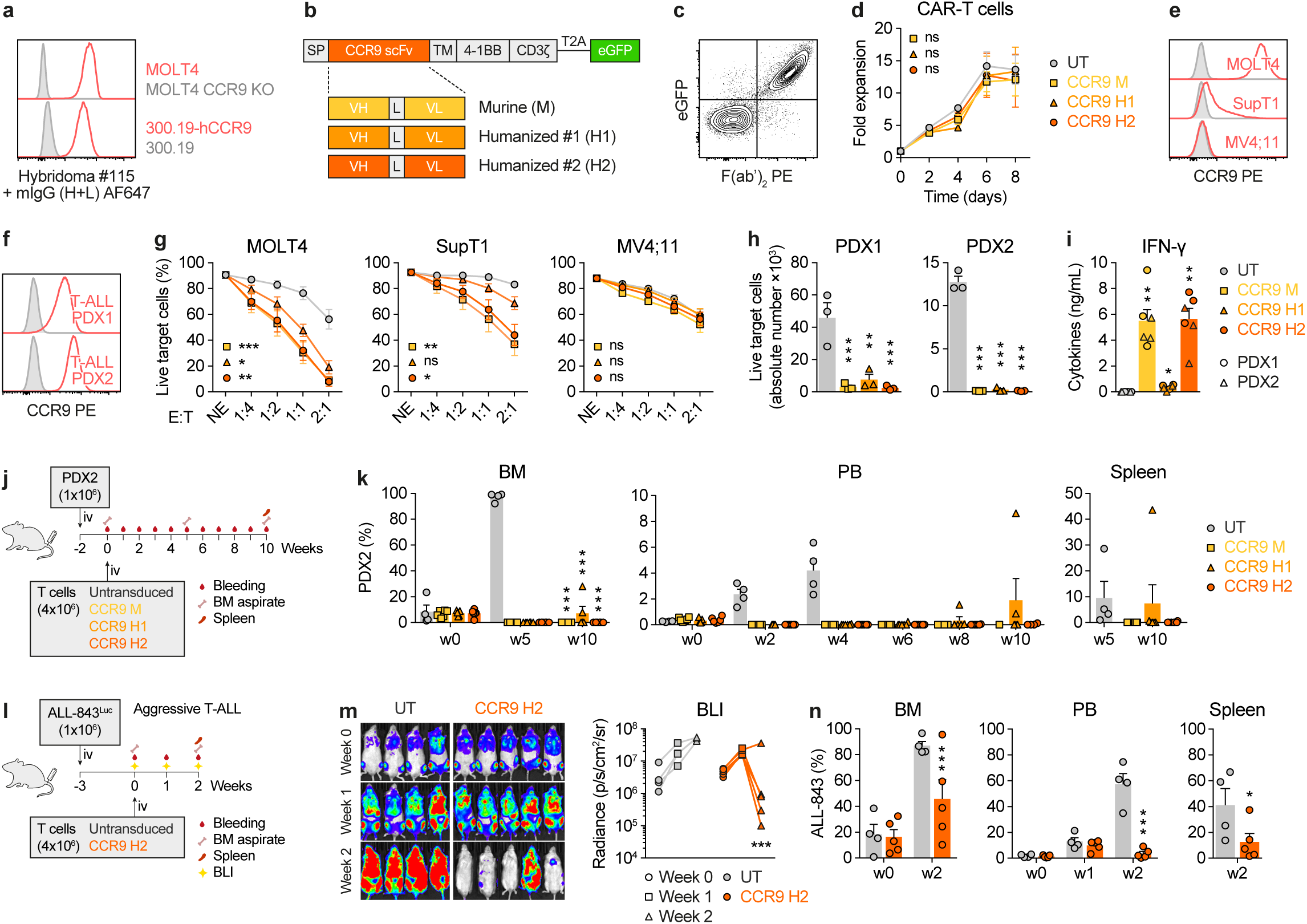
CCR9 CAR-T cells are highly effective against T-ALL. (**a**) Anti-CCR9 hybridoma specifically stains CCR9-expressing cells. (**b**) Cartoon of second-generation CAR constructs. Three scFv versions were generated: scFv derived from the murine (M) hybridoma, and two humanized candidates (H1 and H2). SP, signal peptide; V_H_ and V_L_, heavy and light chains; L, linker; TM, CD8 transmembrane domain. (**c**) Representative flow cytometry plot showing successful detection of transduced CAR-T cells as measured by co-expression of surface F(ab’)_2_ (scFv) and eGFP reporter. (**d**) Proliferation curves for untransduced (UT) and the indicated CCR9 CAR-transduced T cells (*n*=3). (**e**) CCR9 expression in the target T-ALL cell lines MOLT4 (CCR9^high^) and SupT1 (CCR9^dim^) and the CCR9^neg^ AML cell line MV4;11. Isotype control stainings in gray. (**f**) CCR9 expression in two T-ALL PDXs. (**g**) 24 h cytotoxicity mediated by the different murine and humanized CCR9 CAR-T cells against the indicated cell lines at different effector:target (E:T) ratios (*n*=5). NE, no effector T cells. (**h**) Absolute numbers of live target PDX cells after co-culture with untransduced (UT) or the indicated CCR9 CAR-T cells for 24 h at an E:T ratio of 1:1 (*n*=3). (**i**) IFN-γ production by the indicated CCR9 CAR-T cells upon 24 h co-culture with PDXs (E:T 1:1, *n*=3 per PDX). (**j**) *In vivo* experimental design for the assessment of the efficacy of the three indicated CCR9 CAR-T cells against a T-ALL PDX (PDX2, *n*=4-6 mice/group). (**k**) Flow cytometry follow-up of tumor burden in BM, PB, and spleen in the different treatment groups indicated in (j). (**l**) *In vivo* experimental design for the assessment of the efficacy of the selected CCR9 H2 CAR-T cells against a highly aggressive Luciferase-bearing T-ALL PDX (ALL-843, *n*=4-5 mice/group). (**m**) Weekly bioluminescence imaging of mice. Left panel, bioluminescence images. Right panel, bioluminescence quantification. (**n**) Flow cytometry follow-up of tumor burden in BM, PB, and spleen after treatment with UT or CCR9 H2 CAR-T cells. Plots show mean ± SEM.

The T-ALL cell lines MOLT4 and SupT1 (with high and dim expression of CCR9, respectively) and the control (CCR9 negative) acute myeloid leukemia cell line MV4;11, as well as two independent T-ALL PDX samples (**Fig. 2e,f**), were used to assess the *in vitro* cytotoxicity of CCR9 CAR-T cells (**Fig. 2g,h**). Cytotoxic activity was assessed in 24-h co-cultures with untransduced/CAR-T cells at different E:T ratios. CCR9 M and H2 CAR-T cells displayed similar robust and specific cytotoxicity in an antigen density-dependent manner. By contrast, H1 CAR-T cells exhibited slightly inferior killing, particularly with CCR9^dim^ SupT1 cells and, to a lesser extent, with PDX1 blasts (**Fig. 2g,h**). IFN-γ secretion in the co-culture supernatants, used as a surrogate for CAR-T cell activation and cytotoxicity, was highest in CCR9 M and H2 CAR-T cells (**Fig. 2i**).

To test CAR-T cell function *in vivo*, we used two different T-ALL PDX models. In the first model, we compared untransduced and all three CAR-T cells (M, H1, and H2) using a slow growing T-ALL PDX model (PDX2) (**Fig. 2j**). In contrast to untransduced T cells, all three treatment groups were able to control leukemia progression, as assessed by flow cytometry 10-week follow-up in BM, PB, and spleen (**Fig. 2k**, gating strategies in **Fig. S1e**). However, while all mice treated with CCR9 M or H2 CAR-T cells achieved complete response, 2 of 6 mice treated with CCR9 H1 CAR-T cells showed detectable leukemic burden at the endpoint (**Fig. 2k**). This further supports the *in vitro* data demonstrating the higher anti-leukemia efficacy of CCR9 H2 CAR over H1 CAR. Accordingly, CCR9 H2 CAR-T cells were selected for further experiments. To additionally test the robustness of CCR9 H2 CAR-T cells, we used a second, highly aggressive CCR9^+^ luciferase-bearing PDX (ALL-843) (**Fig. 2l**). Bioluminescence follow-up showed disease remission in 4 out of 5 mice (80%) at week 2 post-treatment in the CCR9 H2 CAR-T-treated group, in contrast to disease progression in all control mice (**Fig. 2m**). Disease progression was also monitored by flow cytometric analysis of BM, PB, and spleen, confirming leukemia control in mice treated with CCR9 H2 CAR-T cells but not in control-treated mice (**Fig. 2n**), even in this highly aggressive model. Thus, the humanized CCR9 H2 CAR is highly effective against T-ALL blasts *in vitro* and *in vivo* using several T-ALL cell lines and different PDX models.

### Humanized CCR9 and CD1a dual targeting CAR-T cells for T-ALL

Our group has previously proposed a CD1a-directed CAR for the treatment of patients with CD1a^+^ cortical T-ALL^25,29^. Since both CD1a and CCR9 are safe non-pan-T targets that bypass both fratricide and T cell aplasia, we next immunophenotyped the 180 T-ALL samples for CD1a and CCR9 co-expression (**Fig. 3a**). Using a 20% expression cut-off for each marker, we observed that 51% and 73% of the patients expressed either CD1a or CCR9, respectively (**Fig. 3a**). Remarkably, however, we observed highly heterogeneous leukemic populations for CCR9 and CD1a, with each patient showing a unique co-expression profile, predominating in either CCR9^+^ or CD1a^+^ blasts (**Fig. 3a,b**). Strikingly, the analysis of all patients with >20% expression of either antigen revealed that 86% (155/180) of cases could benefit from dual CCR9/CD1a CAR-T cell therapy.

**Figure 3.**
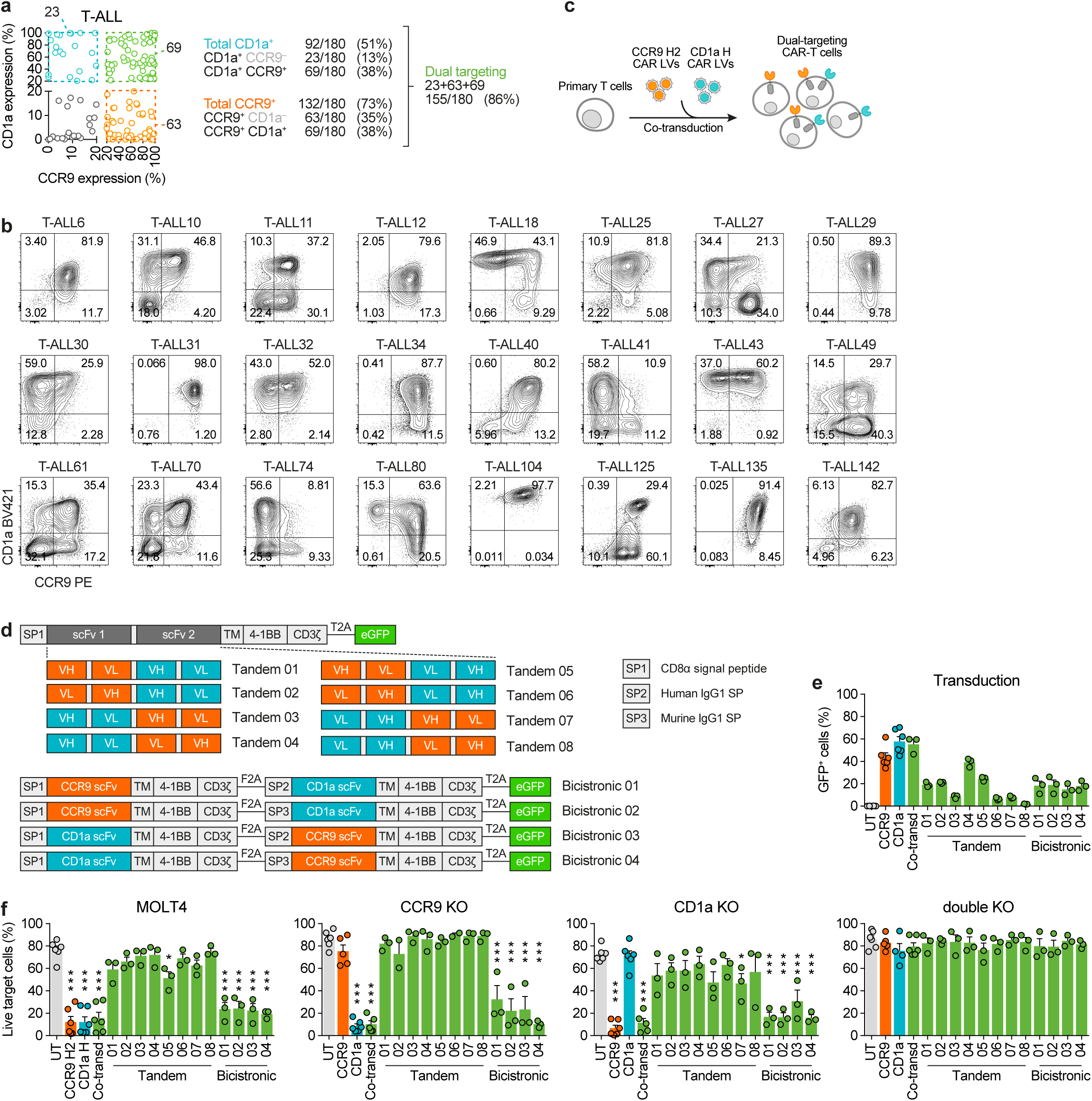
Co-expression of CCR9 and CD1a in T-ALL and molecular strategies for dual targeting with CAR-T cells. (**a**) Flow cytometric analysis of CCR9 and CD1a expression in blasts from 180 T-ALL primary samples, 20% expression cut-off was set to define positivity for each marker. (**b**) CCR9 and CD1a immunophenotypes in 24 representative T-ALL patient samples. Cut-off thresholds for antigen positivity were determined using isotype controls. (**c**) Scheme depicting the generation of dual-targeting CAR-T cells through lentiviral co-transduction of single CARs. (**d**) Cartoons of all CAR constructs tested. Top panels, eight different configurations of tandem CAR constructs with different arrangements of the humanized scFvs (CCR9-CD1a vs. CD1a-CCR9 and V_H_-V_L_ vs. V_L_-V_H_). Bottom panels, four distinct configurations of bicistronic CAR constructs (CCR9-CD1a vs. CD1a-CCR9). Different signal peptides (SP) derived from CD8α (SP1), human IgG1 (SP2), and murine IgG1 (SP3) were used for bicistronic CARs. The CCR9 H2 scFv was used in all versions. (**e**) Transduction efficiencies of single, co-transduced, tandem, and bicistronic CAR-T cells. (**f**) Cytotoxicity assays comparing the specificity and efficiency of the different single, co-transduced, tandem, and bicistronic CAR-T cells against combinatorial phenotypes of MOLT4 T-ALL cells at a 1:1 E:T ratio after 24h of co-culture (*n*=3-6). UT T cells were used as controls. Plots show mean ± SEM.

The malleable nature of many markers has been demonstrated in various subtypes of acute leukemia^40,41^. To gain insight into the clinical-biological impact of the intratumoral phenotypic heterogeneity of CD1a and CCR9, we performed *in vivo* experiments whereby the CD1a^+/–^ and CCR9^−^ leukemic fractions from primary T-ALLs were FACS-sorted and transplanted into NSG immunodeficient mice to evaluate the phenotype of the resulting engraftment (**Fig. S3**). These experiments showed that both CD1a^−^ and CCR9^−^ fractions were capable of engrafting and, importantly, that the graft reproduced the initial leukemia phenotype, where positive and negative populations for CD1a and CCR9 coexist (**Fig. S3**). This marker plasticity suggests that patients with even <20% positive blasts for each marker could benefit from dual treatment (**Fig 3a,b**), thereby increasing not only the number of patients eligible for treatment but also the blast coverage per patient, which would likely contribute to lower immune escape rates.

We therefore generated dual CAR-T cells targeting both CCR9 and CD1a. Several molecular strategies to achieve dual targeting were tested, including eight configurations of tandem CARs (two scFvs in a single CAR molecule), four bicistronic CARs (two independent CAR molecules encoded in one lentiviral vector), and co-transduction with two single CAR-encoding lentiviral vectors simultaneously (**Fig. 3c,d**). A comparison of the transduction efficiency and cytotoxic efficacy for each strategy revealed that the co-transduction strategy achieved significantly higher transduction levels and specific cytotoxic performance using T-ALL cells with combinatorial CCR9/CD1a phenotypes (wt, CCR9 KO, CD1a KO, and double KO) (**Fig. 3e,f**). Thus, co-transduction with single CCR9 CAR and CD1a CAR viral vectors was selected as our dual-targeting CAR-T strategy of choice.

Using combinatorial phenotypes for both antigens of T-ALL cells, we then tested the specificity and efficiency of dual CCR9/CD1a-directed CAR-T cells generated by co-transduction. In contrast to single CAR-T cells, dual CCR9/CD1a CAR-T cells were able to eliminate all target cells in 24-h co-cultures at low E:T ratios as long as one of the antigens remained expressed (**Fig. 4a,b**).

**Figure 4.**
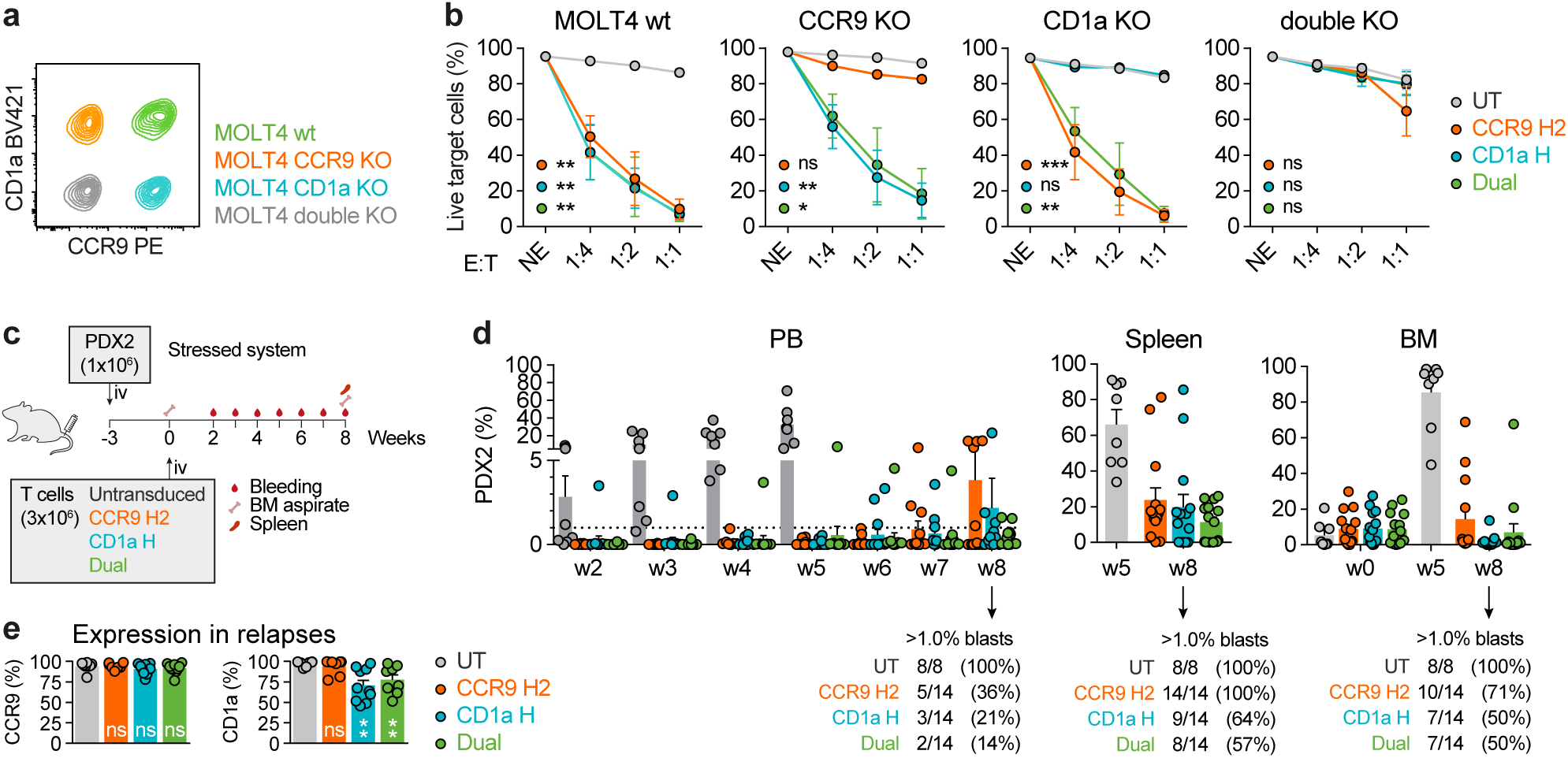
Efficacy of CAR-T cells co-transduced with single CARs. (**a**) Flow cytometry analysis of the MOLT4 cells CRISPR/Cas9-engineered to express combinatorial CCR9/CD1a phenotypes (+/+, –/+, +/–, –/–). (**b**) *In vitro* cytotoxicity assays of the different phenotypes of MOLT4 cells with single CAR (CD1a H or CCR9 H2) T cells or CCR9/CD1a dual-targeting CAR-T cells at different E:T ratios after 24 h of co-culture (*n*=3). (**c**) *In vivo* experimental design for the assessment of the efficacy of CCR9- and CD1a-targeting CAR-T cells against a CCR9^+^CD1a^+^ T-ALL PDX (PDX2) (*n*=8-14 mice/group). (**d**) Flow cytometry follow-up of tumor burden in PB, spleen, and BM after treatment with the indicated CAR treatments. Frequencies of relapsing mice (>1% blasts) for each tissue are indicated. (**e**) Expression of CCR9 and CD1a in CAR-T-resistant blasts. Plots show mean ± SEM.

Next, we tested the *in vivo* efficacy of dual CCR9/CD1a CAR-T cells in a “stress” model against PDX cells expressing both antigens by injecting fewer therapeutic T cells (**Fig. 4c, S1e**). Weekly flow cytometry follow-up in the PB revealed that all CAR-T cell treatments controlled the disease for up to 4-5 weeks. However, when mice were allowed to relapse, the dual CAR-T therapy offered slightly higher rates of complete response (defined as <1% of blasts) in PB and spleen (**Fig. 4d**). In addition, immunophenotyping of the CAR-resistant T-ALL blasts analyzed at the endpoint (week 8) revealed partial downregulation of CD1a in mice treated with CD1a-directed CAR-T cells (both single CD1a or dual CCR9/CD1a CAR-Ts), whereas no downregulation was observed with CCR9 targeting (**Fig. 4e**). Taken together, the immunophenotyping data and the *in vitro* and *in vivo* experimental results support the potential for treating R/R T-ALL with CAR-T cells targeting both CCR9 and CD1a, generated by co-transduction with two single CARs.

### CCR9 and CD1a dual targeting CAR-T cells efficiently eliminate T-ALL with phenotypically heterogeneous leukemic populations

Next, we sought to evaluate the efficiency of co-transduced dual CCR9/CD1a CAR-T cells in the context of phenotypically heterogeneous leukemias. Intratumor phenotypic heterogeneity was recreated by mixing CCR9/CD1a combinatorial phenotypes (CCR9^+^CD1a^+^, CCR9^+^CD1a^−^, and CCR9^−^CD1a^+^) of MOLT4 T-ALL cells in a 1:1:1 ratio (**Fig. 5a**). Time-course cytotoxicity assays revealed complete ablation of the entire leukemic population with dual CAR-T cells, whereas, as expected, single CAR-T cells were unable to eliminate those T-ALL cells that were negative for the corresponding target antigen, leading to leukemic escape (**Fig. 5b**). Identical results were obtained with a MOI of 5+5 and 10+10 for each single CAR vector for the dual strategy in terms of cytotoxicity (**Fig. S4a**), although the number of viral integrations per genome was higher at an MOI of 10+10 (**Fig. S4b**). Similar levels of the pro-inflammatory cytokines IFN-γ, TNF-α, and IL-2 were observed for each treatment (**Fig. 5c**). Finally, we tested the efficacy of dual CCR9/CD1a CAR-T cells in an *in vivo* setting using mixed phenotypes of MOLT4 target cells (**Fig. 5d**). Bioluminescence imaging and BM flow cytometric analysis revealed massive disease control of the highly aggressive heterogeneous MOLT4 cells with respect to both single CAR-T treatments (**Fig. 5e,f**). Taken together, our data highlights the superior efficacy of dual CCR9/CD1a CAR-T cells over single-targeting CAR-T cells in the treatment of T-ALL with phenotypically heterogeneous leukemic populations.

**Figure 5.**
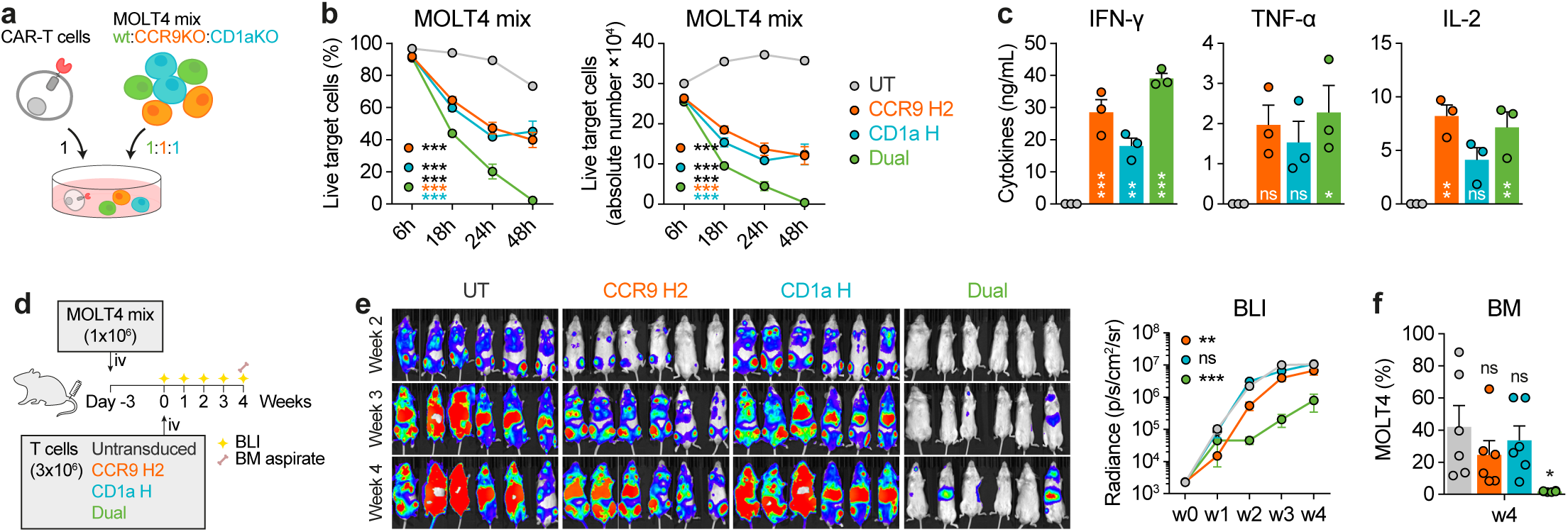
Superior efficacy of dual CCR9 and CD1a CAR-T cells for the treatment of T-ALL with phenotypically heterogeneous leukemic populations. (**a**) Combinatorial phenotypes (CCR9^+^CD1a^+^, CCR9^+^CD1a^−^, CCR9^−^CD1a^+^) of T-ALL cells were mixed at a ratio of 1:1:1 to reproduce phenotypically heterogenous leukemic samples. (**b**) Relative (left) and absolute (right) numbers of live mixed target cells after a time-course cytotoxicity with UT, single CAR (CCR9 H2 or CD1a H) or CCR9/CD1a dual-targeting CAR-T cells at a 1:1 E:T ratio (*n*=3). (**c**) Cytokine production by the indicated CAR-T cells upon 24 h co-culture with target cells (*n*=3). (**d**) *In vivo* experimental design for the assessment of CCR9- and CD1a-targeting dual CAR-T cells against phenotypically heterogeneous Luc-bearing T-ALL target cells (*n*=6 mice/group). (**e**) Weekly bioluminescence imaging of mice (*n*=6 mice/group). Left panel, bioluminescence (BLI) images. Right panel, BLI quantification. (**f**) Flow cytometry analysis of BM tumor burden at the endpoint. Plots show mean ± SEM.

## DISCUSSION

The clinical management of R/R T cell leukemias and lymphomas represents an unmet clinical need. While various immunotherapy strategies such as bispecific antibodies and CAR-T cells have revolutionized the treatment of B cell leukemias, lymphomas, and multiple myeloma, with several products approved by the FDA/EMA, these immunotherapies are much less advanced and have not been approved for T cell malignancies^11^. The primary challenge for the implementation of adoptive immunotherapies in T cell malignancies is the shared expression of surface membrane antigens between tumoral cells and healthy/non-leukemic T cells. This implies that the expression of a CAR targeting any pan-T antigen would most likely result in toxicities such as CAR-T fratricide and T cell aplasia^11,42^. Additionally, the shared expression of antigens between effector and tumor T cells may complicate the manufacturing process of autologous T cell therapies due to blast contamination of the leukapheresis products^10^. Clinical trials with CD7 CAR-T cells often employ stringent inclusion criteria, including a maximum allowable blast threshold (NCT06064903). This threshold is established to ensure that the CAR-T cell production is free of tumor cells, thereby preventing accidental CAR transduction of tumoral T cells and avoiding potential interference from blasts during the activation and expansion phases of the CAR-T cell product.

To overcome these drawbacks, the current trend is to use allogeneic T lymphocytes to bypass potential blast contamination. However, this strategy requires multiple CRISPR/Cas9-mediated gene edits to eliminate molecules such as the target antigen and the T cell receptor (TCR), thereby preventing fratricide and GvHD^15,16^. This strategy is only feasible with “off-the-shelf” effector cells, as the technical and regulatory complexity of CRISPR/Cas9-mediated genome editing makes it difficult to implement with autologous T cells derived from patients in critical clinical states. Crucially, previous studies have demonstrated the negative impact of TCR elimination and genomic manipulation of T cells on the persistence of CAR-T cells and their genomic/chromosomal stability^43,44^. This point is very important in light of recent clinical studies and subsequent FDA investigations into the occurrence of neoplasms secondary to CAR-T therapy^45–49^.

To circumvent the limitations of adoptive cell therapies for T cell malignancies, it would be ideal to redirect effector cells against non-pan-T targets present in the tumor but absent in healthy tissues. This approach facilitates the manufacture of autologous CAR-T cells and avoids both fratricide and immune toxicity^23–28^. In this context, we previously identified CD1a as a *bona fide* immunotherapeutic target for the treatment of cortical T-ALL with a safe profile in non-hematopoietic and hematopoietic tissues^25,29^. This led us to generate and preclinically validate CD1a-directed CAR-T cells, which are now being tested in a phase I clinical trial (NCT05679895). However, CD1a only covers cases of cortical T-ALL, a subtype that accounts for ∼30% of all diagnosed T-ALL cases, while sparing other T-ALL subtypes associated with higher refractoriness and relapse rates^1,27,50,51^. Here, we identify CCR9 as a target expressed in ∼72% of diagnostic T-ALL cases and, importantly, in ∼92% of relapses. Of note, Maciocia and colleagues previously proposed CCR9 as a target for T-ALL and elegantly reported similar expression and safety data^27^. Importantly, these expression rates are maintained in subtypes with poor prognosis and very high rates of refractoriness and relapse, such as early T cell progenitor ALL, an entity with unmet clinical need^52^. Crucially, CCR9 is not expressed on normal circulating T cells or other hematopoietic or non-hematopoietic tissues, with the exception of a subset of B cells, thymocytes, and a small fraction of intestinal-resident lymphocytes. This was expected, as CCR9 expression plays a key role in the homing of specific immune cells to the thymus and small intestine^53–55^. Although it has not been reported that CAR-T cells can reach the thymus, this could be of great therapeutic benefit in the case of CCR9 CAR T cells given that many pre-leukemic clones and leukemic-initiating cells in T-ALL are present at very early developmental stages^56^. Furthermore, many studies in non-oncological pediatric and adult patients who have undergone thymectomy have demonstrated immune memory and a complete T cell repertoire^57,58^. In addition, the CCL25-CCR9 axis has been shown to play a role in inflammatory bowel disease, and previous clinical trials have used CCR9 small molecule inhibitors with no reported serious therapy-related toxicities, further supporting the safety of CCR9 as a therapeutic target^59–62^. Collectively, these data support CCR9 as a safe target with promising potential for a large proportion of R/R T-ALL cases.

A major strength of our work is the immunophenotypic characterization of CCR9 and CD1a expression in a cohort of 180 primary T-ALL cases. This analysis revealed the existence of significant intratumoral phenotypic heterogeneity, with double-positive, single-positive, and double-negative fractions coexisting within the same sample. This suggests that dual CAR-T cell therapy targeting both antigens expressed in heterogeneous leukemias, as is the case in a large percentage of R/R T-ALL cases, would increase the number of patients eligible for treatment, provide greater blast coverage, and potentially reduce the likelihood of phenotype/antigen escape. In addition, transplantation of CCR9- and CD1a-negative fractions into immunodeficient mice demonstrated that leukemic engraftment reproduces the phenotypic heterogeneity of the original leukemia regardless of the input. This provides clear evidence of the plasticity of these antigens and extends the applicability of such dual-targeting immunotherapy to patients without the need for high antigen positivity rates to achieve deep responses. Overall, these results highlight the benefit of a dual-targeting strategy even in patients with leukemic populations that are only partially positive for one of the markers.

Based on these clinico-biological data, we generated a CCR9-specific hybridoma and derived the scFv sequence to generate a second-generation CAR. The murine scFv was humanized by CDR grafting, and one of the humanized CAR candidates (H2) proved to be as potent as the murine CAR and was ultimately selected to minimize potential immunogenicity in humans. We next leveraged our humanized CD1a H CAR and CCR9 H2 CAR to generate dual-targeting strategies. Of the various possible molecular strategies to direct T cells against two molecules^63^, we focused on three: co-transduction with two lentiviral vectors each encoding a different single-targeting CAR, multiple configurations of tandem CARs (two scFvs within the same CAR molecule), and bicistronic CARs (two separate CAR molecules with different specificities encoded in one lentiviral vector). Despite our previous experience in generating tandem CARs targeting other antigens^36^, none of the possible tandem CAR configurations worked, possibly due to the biochemical properties and steric hindrance associated with these specific scFvs, or perhaps due to the nature and mechanisms of recognition of the receptors, where CCR9 is largely embedded in the cell membrane and CD1a has a fully exposed and large ectodomain that makes the tandem CAR configuration unsuitable. Both co-transduction and bicistronic CAR strategies showed efficacy, but we chose co-transduction for further study due to the higher rates of transduction achieved. We then demonstrated the functional advantage of dual-targeting CAR-T cells generated by co-transduction with CCR9- and CD1a-directed CARs over single-targeting CAR-T cells for the treatment of phenotypically heterogeneous T-ALL cases. Our study provides an exhaustive molecular comparative study of all dual-targeting CAR-T cell strategies, confirms that the development of dual strategies is not trivial, and establishes a unique foundation and knowledge for the applicability of our strategy and that of future constructs.

In conclusion, the proposed CAR-T cell strategy targeting two non-pan-T antigens that are absent in normal T cells and barely expressed in other healthy tissues may achieve a large blast coverage and benefit a very significant proportion of R/R T-ALL patients, while preventing T cell fratricide and aplasia, and will obviate the need for regulatory-challenging genome engineering approaches and alloHSCT in patients after CAR-T therapy to rescue T cell aplasia. The fact that CCR9 is also highly expressed in several subtypes of solid tumors with poor prognosis^64–71^, together with the fact that CCR9 is the only canonical receptor of the chemokine CCL25, opens up enormous possibilities for the adoptive immunotherapy of cancers beyond T-ALL using either antibody scFv-based or CCL25 zetakine-based CARs^72,73^.

## ACKNOWLEDGMENTS AND FUNDING

The authors wish to acknowledge Virginia C Rodríguez-Cortez, Pau Ximeno-Parpal, Carla Panisello, Ángela Meseguer Girón, Patrizio Panelli, Alex Bataller, and Patricia Fuentes for their technical support.

Research in PM laboratory is supported by CERCA/Generalitat de Catalunya and Fundació Josep Carreras-Obra Social la Caixa for core support, the European Research Council grants (ERC-PoC-957466 IT4B-TALL, ERC-PoC-101100665 BiTE-CAR); H2020 (101057250-CANCERNA), the Spanish Research Agency (AEI) (PID2022-142966OB-I00 funded by MCIN/AEI/10.13039/501100011033 and FEDER Funds, and CPP2021-008508, CPP2021-008676, CPP2022-009759 funded by MICIU/AEI/10.13039/501100011033 and the European Union NextGenerationEU/PRTR); “Ayudas Merck de Investigación” from Health Merck Foundation, and grant PID2022-136554OA-I00 funded by MICIU/AEI 10.13039/501100011033 and the European Regional Development Fund (ERDF)/EU to DS-M; the AEI grant PID2022-138880OB-I00 to MLT, the Spanish Association Against Cancer (AECC, PRYGN234975MENE, PRYGN211192BUEN), the Uno Entre Cien Mil Foundation to MLT and to CB, and the ISCIII-RICORS within the Next Generation EU program (plan de Recuperación, Transformación y Resilencia). AF was supported by a Juan de la Cierva postdoctoral fellowship (FJC2021-046789-I), VMD by a Torres Quevedo contract (PTQ2020-011056), AG-P by an industrial PhD fellowship (DIN2022-012556) and NT by a PhD fellowship (FPU19/00039) from the Spanish Ministry of Science and Innovation.

## AUTHOR CONTRIBUTION

Conception and design of the study: NT, DS-M, PM; sample acquisition of data: NT, MJM, AM-M, JA, MG-P, HR-H, NF-F, AG-P, AF, TV-H, MV, CB, PE, EAG; analysis and interpretation of data: NT, MJM, NF-F, MG-M, PE, VMD, MLT, DS-M, PM; sample preparation and clinical data: BV, IJ, AC-E, AB, HC, EG, JR, MD-B, MR-O, MT; writing: NT, DS-M, PM; review and editing: all authors; guarantors: PM.

---

**Suppl Fig 1.**
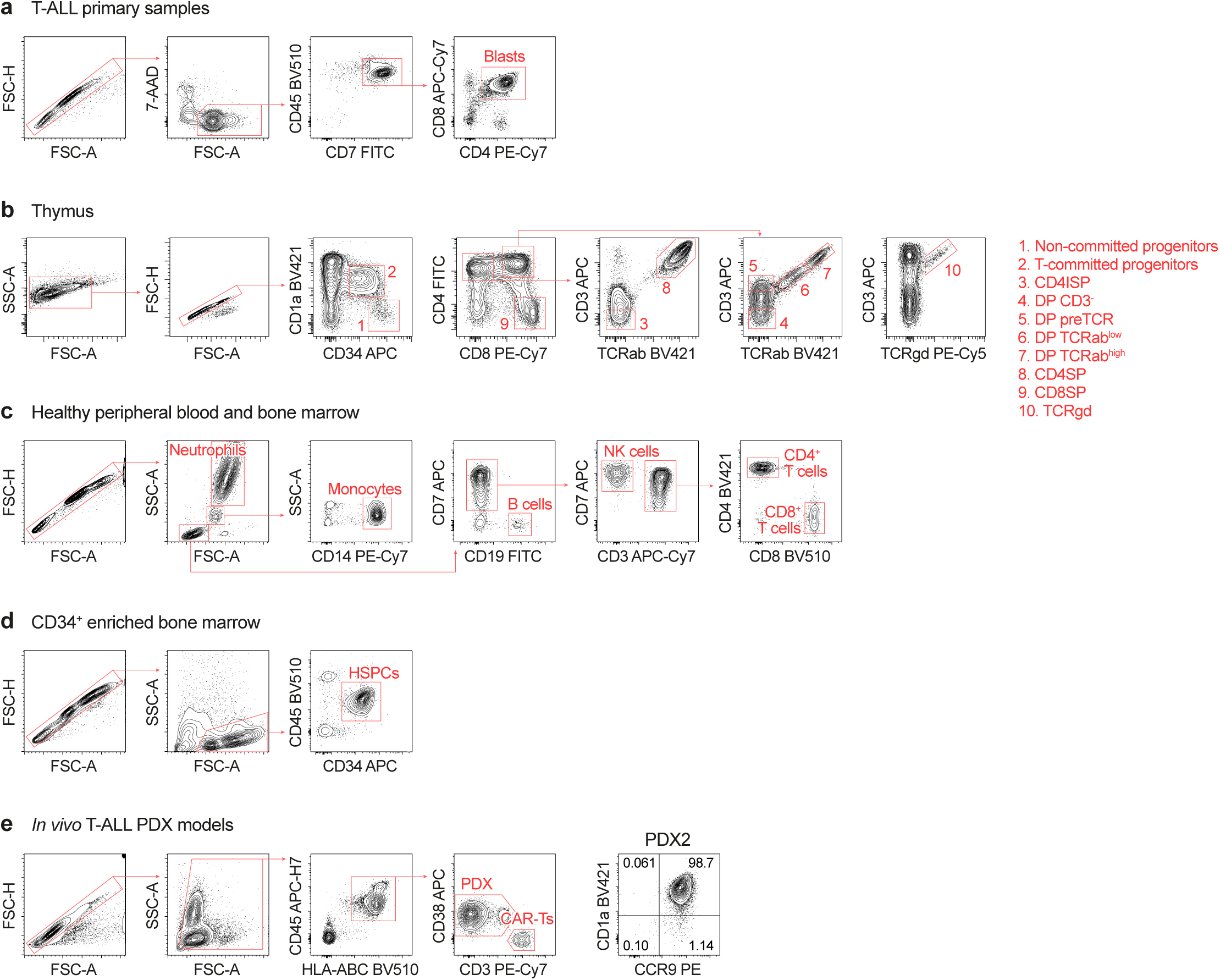
Gating strategies for flow cytometry analyses. (**a**) Flow cytometry gating strategies for T-ALL primary samples. Blasts were identified as CD7^+^CD45^dim^ and were further gated based on CD4 and CD8 expression to exclude healthy (single CD4^+^ and single CD8^+^) T cells from the analysis. (**b-d**) Flow cytometry gating strategies for the indicated cell populations within healthy thymus (**b**), PB and total BM (**c**), and MACS-enriched BM CD34^+^ HSPCs (**d**). (**e**) Flow cytometry gating strategies for *in vivo* PDX assays. A representative phenotype of PDX2 is included.

**Suppl Fig 2.**
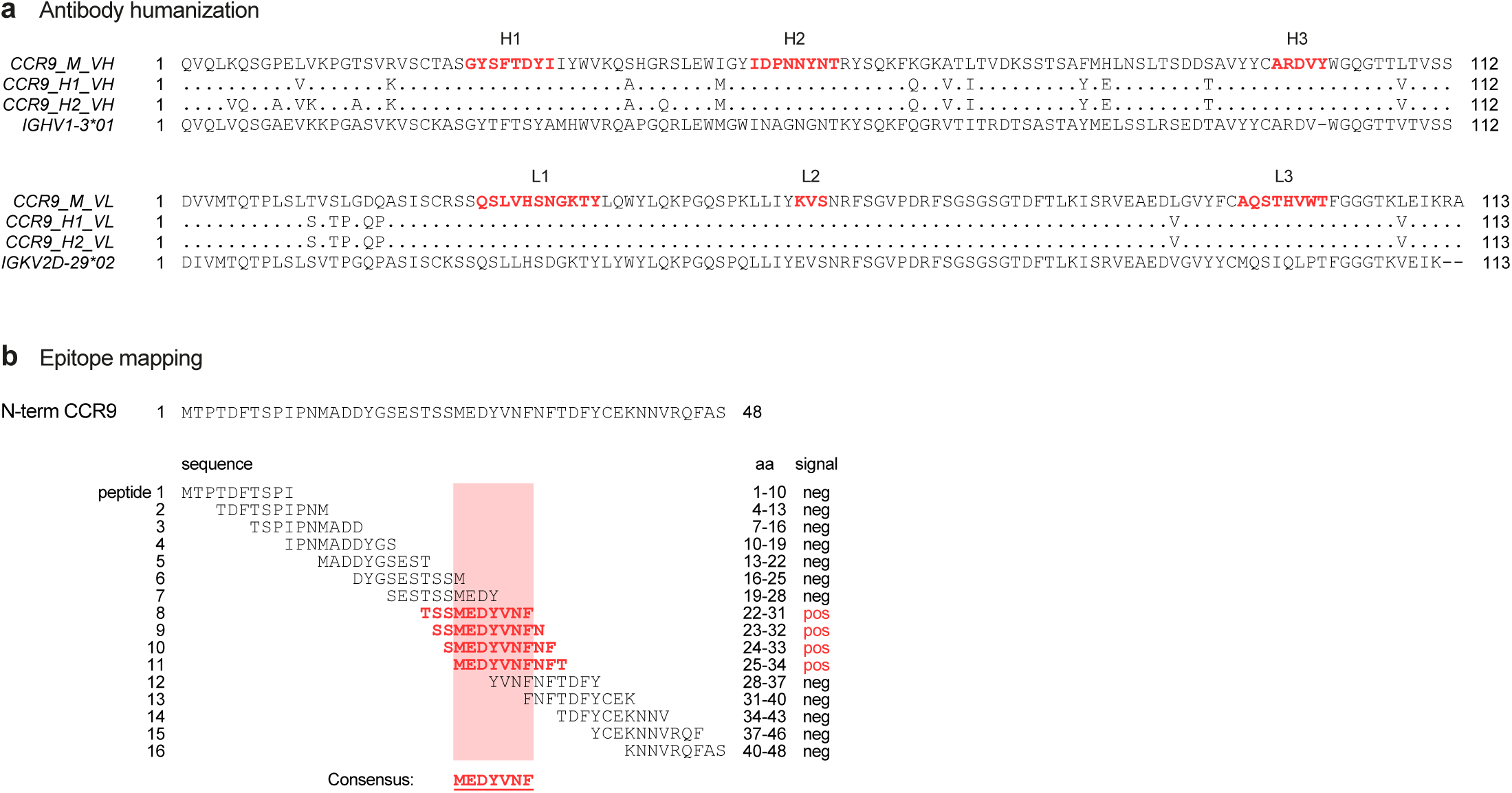
CCR9 binder characterization. (**a**) Protein sequence alignment of the original murine heavy (V_H_) and light (V_L_) chains with their humanized counterparts (H1 and H2) and the most structurally similar human immunoglobulin gene. CDRs are highlighted in red, and only mutated residues in the humanized sequences are shown. (**b**) Epitope mapping of the anti-CCR9 binder. Overlapping peptides were derived from the extracellular N-terminus tail of CCR9 and tested for binding to recombinant anti-CCR9 H2 scFv by ELISA (ProteoGenix). The consensus sequence is indicated.

**Suppl Fig 3.**
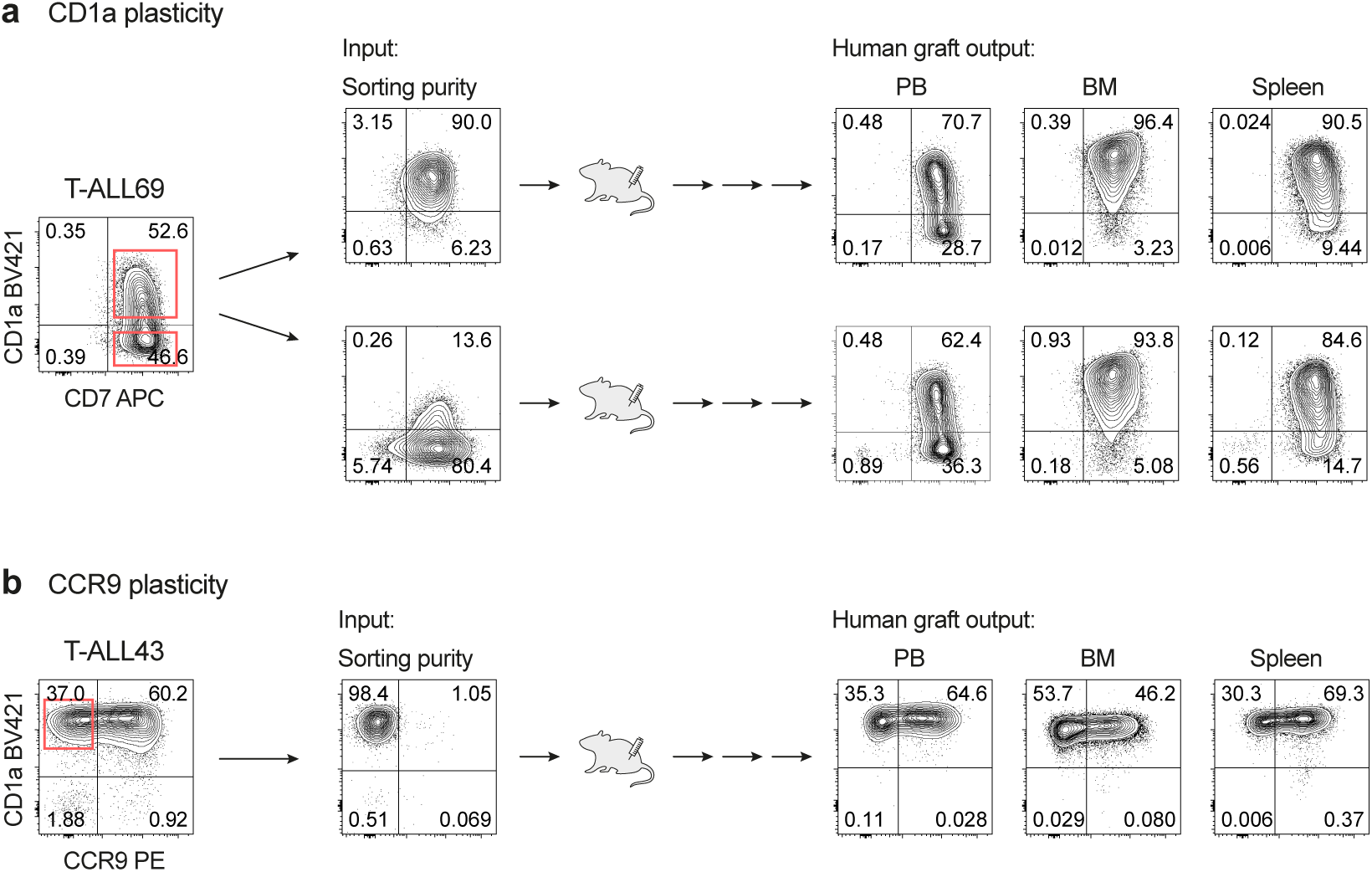
Expression plasticity of CD1a and CCR9. Two primary T-ALL samples with variable expression of CD1a (**a**) and CCR9 (**b**) were sorted, and the purified CD1a^+/–^ or CCR9^−^ fractions (purity: 81-98%) were transplanted into NSG mice. Leukemic grafts were followed up biweekly and mice were sacrificed for leukemia immunophenotyping upon graft detection in PB.

**Suppl Fig 4.**
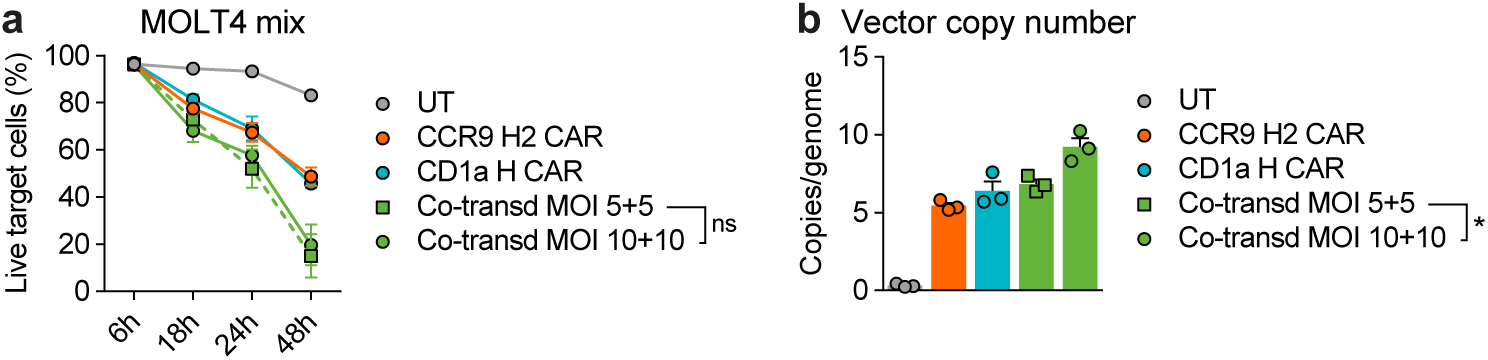
Impact of MOI in co-transduced dual CAR-T cell assays. (**a**) Time-course cytotoxicity comparing co-transduced dual CAR-T cells using a MOI of 5 *versus* 10 for each CAR against mixed target cells at a 1:1 E:T ratio (*n*=3), as described in Fig. 5a. (**b**) Vector copy number (VCN) in co-transduced dual CAR-T cells using a MOI of 5 *versus* 10 for each CAR (*n*=3).

